# Decomposing Parts of Speech: Cerebral Encoding of Semantic and Grammatical Dimensions and Their Combinational Structure

**DOI:** 10.64898/2026.10.29.761822

**Authors:** Zirui Huang, Natalia Bekemeier, Alexis Hervais-Adelman

## Abstract

Conventional part-of-speech (PoS) labels such as nouns, verbs and adjectives conflate lexical-semantic roots, grammatical functions, and their combinations, making it unclear whether neural sensitivity to PoS reflects either dimension or the way they are combined. We adopted Croft’s typological framework to decompose major PoS classes into semantic root type and grammatical function and define prototypical nouns, verbs, and adjectives through the prototypical mappings between semantic root and grammatical function. We fitted time-resolved encoding models to magnetoencephalography (MEG) data from three participants who each listened to approximately ten hours of narrative speech. We examined the unique predictive contributions of semantic root, grammatical function, their interaction, and a binary prototypicality predictor, and whether the three prototypes exhibited distinct neural response dynamics.

Semantic root and grammatical function contributed independently to neural responses during naturalistic language comprehension, with additional contributions from their interaction beyond the main effects. Root effects were predominantly left-lateralized, function effects were more bilateral, and interaction effects were concentrated in the left temporal and temporoparietal cortex, with overlap in left temporal regions. Further analyses revealed significant neural sensitivity to the distinction between prototypical and nonprototypical combinations in two of the three datasets. Prototypical nouns and verbs showed distinct coefficient time courses in all three participants, most consistently in left superior temporal cortex, whereas adjective-related contrasts were less consistent. These within-participant findings support a decompositional account of neural word-class sensitivity involving semantic, grammatical, and the mappings between them within distributed, partially overlapping cortical networks.

## Introduction

Parts of speech (PoS) are fundamental to linguistic theory because they provide a basic word categorization system for describing how words behave and combine in sentences. Understanding neural sensitivity to PoS is therefore important for explaining how the brain represents lexical categories and uses category information during language comprehension. Yet traditional PoS labels conflate several potentially separable linguistic dimensions into a single category: the lexical-semantic root, the grammatical function a word serves in a context, and how root and function are combined. For the core PoS categories of nouns, verbs, and adjectives, their semantic roots can broadly denote object, action, or property, while their grammatical functions can be reference, modification, and predication. However, semantic roots and grammatical functions do not map onto each other in a one-to-one manner. For example, a word with an action-denoting semantic root like “sleep” can serve a predication function in “she is sleeping” or a modification function in “a sleeping baby”. Different combinations of semantic roots and grammatical functions can thus give rise to different linguistic operations to derive the intended PoS categories, with some root-function combinations more prototypical than others. Consequently, the observed neural sensitivity to core PoS categories, namely, nouns, verbs and adjectives, is inherently ambiguous: underlyingly it may reflect semantic content, grammatical function, or their particular combinations. A decompositional approach to PoS is therefore needed to disentangle these contributions while also examining whether and when prototypical nouns, verbs and adjectives exhibit distinct neural response patterns.

Neural sensitivity to PoS categories has been widely reported (Damasio & Tranel, 1993; Pulvermüller et al., 1999; Federmeier et al., 2000; Shapiro et al., 2006). However, the linguistic basis of these effects remains debated, as it is often unclear whether they are driven by their grammatical class or by semantic meanings. Vigliocco and colleagues focused on noun-verb distinction and directly examined this ambiguity by manipulating semantic and grammatical factors independently. They found neural activity was primarily modulated by sensory and motor semantic properties rather than grammatical class (Vigliocco et al., 2006). Similarly, when object-action semantics were controlled in an event-naming task, differences between action nouns and verbs reflected the demands of producing different word forms rather than a stable noun-verb distinction (Siri et al., 2008). Later in a review, Vigliocco et al. (2011) argued that grammatical class alone is insufficient to explain the clear neural separability between nouns and verbs. Instead, word class effects emerge from semantic–pragmatic and distributional information and become more prominent when morphosyntactic or sentence-level integration is required. These studies highlight the need to distinguish semantic and grammatical information when interpreting PoS-related neural effects. However, previous research has largely examined the semantic and grammatical sources of PoS-related neural effects through the noun-verb distinction, and asked whether semantic content or grammatical class better accounts for the noun-verb distinction, without explicitly representing semantic root type and grammatical function as dimensions whose independent and joint contributions could be tested.

To distinguish how different linguistic dimensions contribute to PoS-related neural effects, we adopted Croft’s typological framework, which provides a principled way to separate semantic root type from grammatical function (Croft, 2001, 2002). The framework classifies three semantic root types: objects, properties, and actions, alongside three grammatical functions: reference, modification, and predication (Table 1). Importantly, Croft’s framework goes beyond separating semantic and grammatical information by proposing three prototypical mappings: object × reference, action × predication, and property × modification, corresponding to prototypical nouns, verbs, and adjectives, respectively. These prototypes are grounded in recurring cross-linguistic patterns of how semantic concepts are used in grammatical constructions, providing a common basis for defining and comparing word classes across languages (Croft, 2001; Kemmerer, 2014). The remaining six combinations represent nonprototypical mappings in which semantic root types serve other non-prototypical grammatical functions. This framework therefore allows us not only to examine the independent and combinational contributions of semantic root and grammatical function, but also to address a further question: whether the three theoretically defined word-class prototypes exhibit distinct neural response patterns, and if so, where and when these differences emerge.

**Table 1.** Decomposition of core content word classes in Croft’s framework. Rows represent three semantic root types, namely, object, property and action. Columns represent three grammatical functions, namely, reference, modification, and predication. The highlighted diagonal cells indicate the prototypical root-function combinations: object–reference, property–modification, and action–predication, corresponding prototypically to nouns, adjectives, and verbs in English. The off-diagonal cells illustrate nonprototypical combinations in which the same type of semantic Root serves a different grammatical Function.

|  |  | *Grammatical Function* |  |  |
| --- | --- | --- | --- | --- |
|  |  | reference | modification | predication |
| *Lexical-semantic concept* | object-roots | *Visit a **school**.* | ***School** bus* | *She is well-**schooled**.* |
|  | property-roots | *The **redness** of her cheeks.* | ***red** flower* | *Rose is **red**.* |
|  | action-roots | *Get some **sleep**.* | ***Sleeping** girl* | *She **sleeps** well.* |

We applied Croft’s framework to annotate words in the narrative *The Adventures of Sherlock Holmes*, which has been used as naturalistic speech comprehension material for a publicly available 10-hour within-participant MEG dataset by Armeni et al. (2022). The stimulus materials consisted of approximately 10 hours of speech, with around 85,000 words. Three participants each listened to one hour of narrative speech each day over ten recording sessions. This intensive within-participant design supports the estimation of time-resolved encoding models separately for each participant. We employed time-resolved encoding modelling using a regression-based event-related field (ERF) framework (Heilbron et al., 2022; Smith & Kutas, 2015a, 2015b; Crosse et al., 2016) to predict neural responses based on linguistic predictors of interest, including Croft-decomposed parameters: semantic root, grammatical function and their combinations. This approach allows us to determine (i) whether semantic root and grammatical function make independent contributions to neural responses during naturalistic language comprehension; (ii) whether the combinational structure of semantic root and grammatical function contributes additionally to their individual effects; and (iii) whether and when the three prototypical combinations underlying nouns, verbs, and adjectives exhibit distinct neural response dynamics.

## Results

### Semantic root and grammatical function preserve core parts-of-speech information

We first evaluate the overall predictive performance of the theoretically motivated Croft decomposition, asking whether Croft-based models explained as much or more variance in neural responses as a model using conventional PoS labels. We constructed three regression-based ERF models that were otherwise identical: a model containing conventional part-of-speech labels generated through automated tagging by spaCy (Honnibal et al., 2020) at the content-word level (hereafter, spaCy PoS), a model containing semantic root and grammatical function labels as predictors at content word level (hereafter, the Croft additive), and a model that additionally contained the semantic root and grammatical function interaction labels at the content word level (hereafter, the Croft interactive). All three models contain the same control linguistic regressors: lexical surprisal and word frequency for each word; semantic mismatch and contextual dissimilarity for each content word; phoneme surprisal, uniqueness point, and acoustic parameters including broadband amplitude envelope variance, pitch, and spectral variation. Prediction performance was quantified as the cross-validated correlation between predicted and observed source activity, averaged over parcels in the language-network mask (See Supplementary Figure 1).

The Croft additive model modestly but reliably outperformed the conventional spaCy part-of-speech model in all three datasets (participant 1, Δr = 1.47 × 10⁻⁴, Holm-corrected p = 0.017; participant 2, Δr = 2.43 × 10⁻⁴, Holm-corrected p < .001; participant 3, Δr = 2.21 × 10⁻⁴, Holm-corrected p = 0.003), see Figure 1. Adding the semantic root and grammatical function interaction did not produce a further reliable increase over the Croft additive model (participant 1, Δr = -7.30 × 10⁻^5^, Holm-corrected p = 0.055; participant 2, Δr = -5.50 × 10⁻^5^, Holm-corrected p = 0.122; participant 3, Δr = 2.96 × 10⁻^5^, Holm-corrected p = .939). Relative to the conventional spaCy PoS model, the interaction model performed reliably better for participants 2 and 3 (participant 2, Δr = 1.88× 10⁻^4,^ Holm-corrected p = 0.003; participant 3, Δr = 2.51 × 10⁻⁴, Holm-corrected p = 0.015), but not for participant 1 (participant 1, Δr = 7.4 × 10⁻^5^, Holm-corrected p = 0.267).

**Figure 1.**
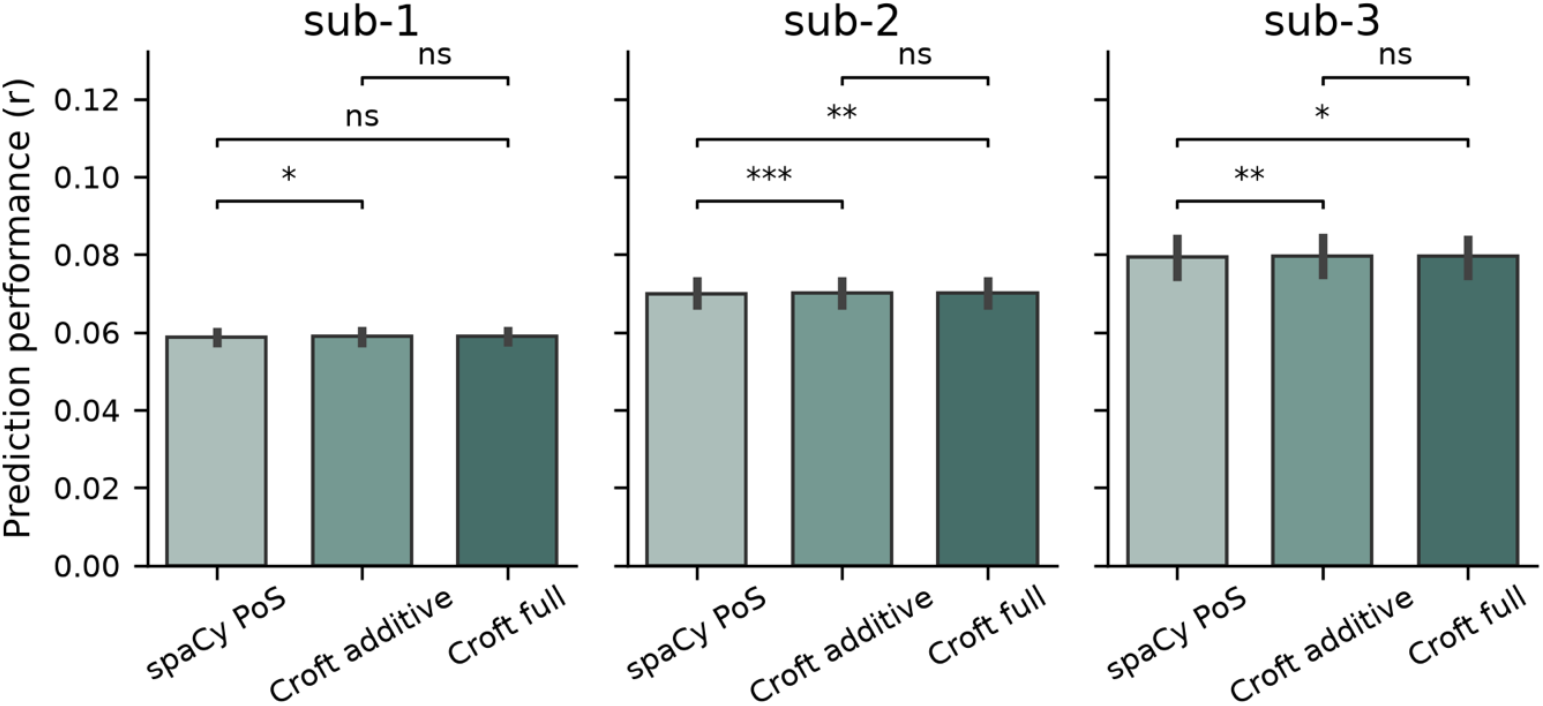
Comparison of overall neural prediction performance between conventional PoS and Croft-based models. Bars show the mean cross-validated correlation (r) between predicted and observed MEG responses for each participant. Prediction performance was first averaged across predefined language-related parcels within each held-out fold and then averaged across folds; error bars indicate 95% confidence intervals across folds. The spaCy PoS model represented content words using conventional PoS categories; the Croft additive model included the main effects of semantic root and grammatical function; and the Croft full model additionally included their interaction. All three models contained the same acoustic, lexical, and contextual control predictors. Statistical annotations indicate paired comparisons of fold-wise prediction performance, p-values were Holm-corrected across the three comparisons separately for each participant.

### Semantic root and grammatical function make distinct predictive contributions

After confirming that the theoretically motivated Croft decomposition does not come at the expense of predictive performance, we then assessed the unique contribution of semantic root and grammatical function in predicting brain responses, by comparing the Croft additive model with permuted models in which the labels of the predictor of interest were permuted while the model structure and all other predictors were retained. Unique contribution was quantified as ΔR², the difference in explained variance between the true-label and permuted-label models. Statistical inference was performed separately within each participant dataset using threshold-free cluster enhancement (TFCE) with correction over source parcels.

We found that permuting semantic root labels significantly reduced prediction accuracy across spatially distributed cortical regions in all three subjects (Figure 2a). Nineteen significant parcels overlapped across three subjects, and this overlap was strongly left-lateralized (17 left, 2 right), predominantly in the left temporal cortex, with additional involvement of the left anterior frontal and right temporal cortices. Unique contributions of grammatical function were even more broadly distributed in the cortical regions across all three subjects (Figure 2c), with more shared significant parcels across subjects (41 parcels). Unlike the effects of semantic root, the grammatical function overlap was bilateral (21 left, 20 right), extending across both frontal and temporal cortices.

**Figure 2.**
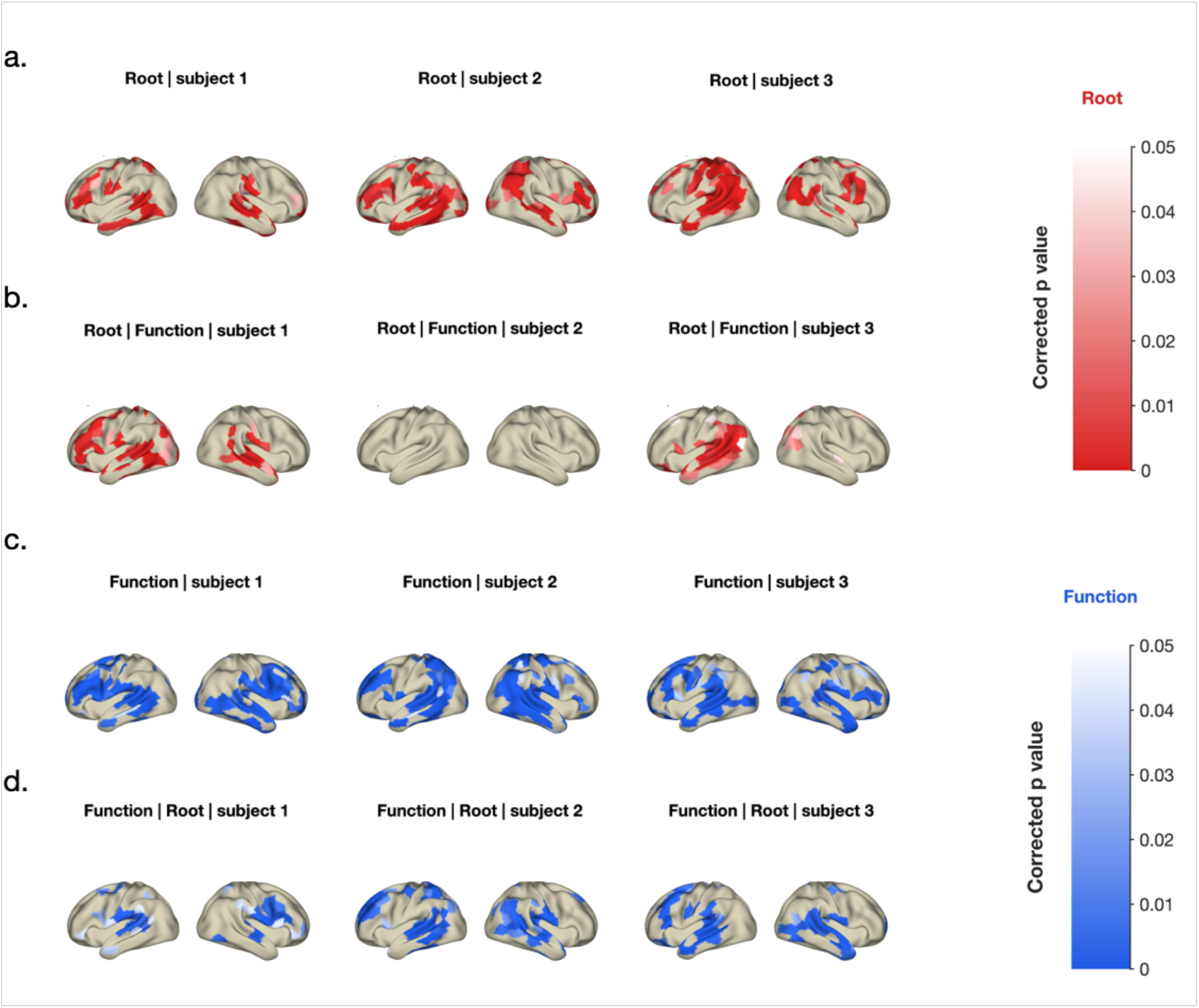
Spatial distribution of the unique contribution of predictors of interest to neural prediction. Inflated cortical surface maps show the parcel-wise unique contribution of each predictor separately for each participant. The unique contribution was quantified as the difference in cross-validated explained variance (ΔR²) between the Croft model containing the original annotations for the predictor of interest and a matched-complexity control model in which those annotations were permuted while the model structure and number of predictors were preserved. The colour scale represents TFCE-corrected p-values, with darker colours indicating smaller p-values. Parcels with p ≥ .05 are transparent. **a.** Unique contribution of semantic root annotations in the Croft additive model. **b.** Unique contribution of semantic root after controlling for their grammatical functions. **c.** Unique contribution of grammatical function annotations in the Croft additive model. **d.** Unique contribution of grammatical function after controlling for their semantic roots.

Since certain semantic root-grammatical function combinations occur more frequently than others in natural language, e.g., words with “object” semantic root are more likely to have a grammatical function of “reference”, the contributions of semantic root and grammatical function might be difficult to disentangle. To account for this statistical dependency, we also used conditional permutation to further determine whether the unique contribution of each dimension remains informative after accounting for the other. For semantic root, we constructed permuted models in which the semantic root labels were shuffled only among words sharing the same grammatical function. For grammatical function, vice versa. This procedure preserves the statistical association between the two dimensions while disrupting the correspondence between the permuted labels and the neural responses. We found that when root was permuted conditionally on function, the unique contribution of semantic root was still preserved for subject 1 and subject 3 with very similar spatial distribution (Figure 2b), but for subject 2, the variance explained by semantic root was not significant after multiple comparisons correction. In contrast, the conditional grammatical function effect remained significant across all three subjects (Figure 2d), with 16 parcels shared across all three datasets (8 left, 8 right) mostly in bilateral temporal cortices.

### Spatially localized sensitivity to combinations of root and function

Although adding the semantic root and grammatical function interaction did not reliably improve overall prediction performance when averaged across the language network, this does not rule out its potential contributions in specific cortical regions. We therefore examined the unique contributions of the semantic root and grammatical interaction at the parcel level by comparing an unpermuted interaction model with a matched model containing randomized interaction regressors, to see if the interaction provides reliable additional information in predicting neural responses.

We found consistent and extended significant contributions of the interactions in addition to the contributions of semantic root and grammatical function across all three subjects (Figure 3a). 27 parcels were shared across all three subjects, predominantly in the left hemisphere (22 left, 5 right), concentrated in left temporal and temporoparietal regions.

**Figure 3.**
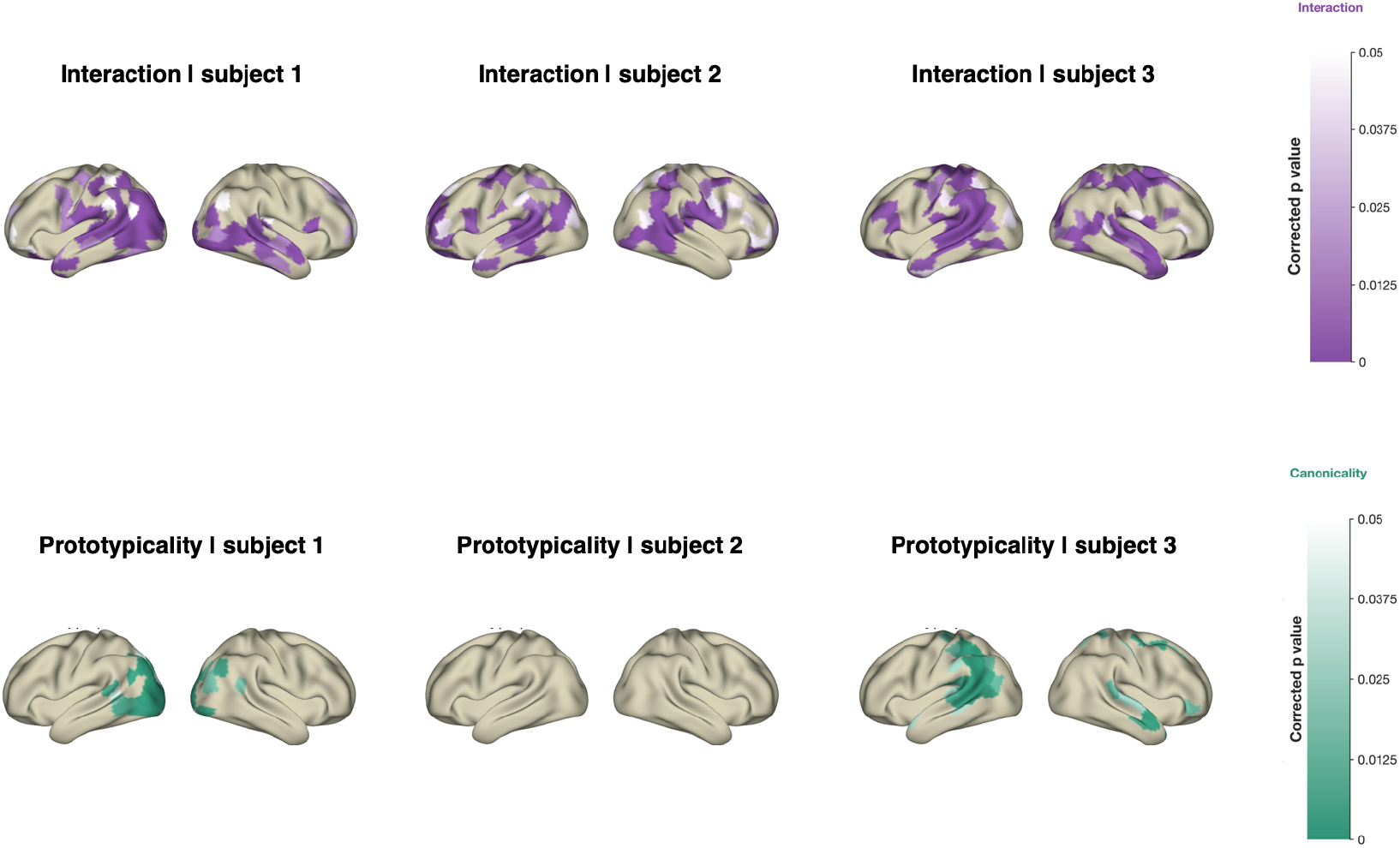
Spatial distribution of semantic root and grammatical interaction. Inflated cortical surface maps show the parcel-wise unique contribution of each predictor separately for each participant. The unique contribution was quantified as the difference in cross-validated explained variance (ΔR²) between the Croft model containing the original annotations for the predictor of interest and a matched-complexity control model in which those annotations were permuted while the model structure and number of predictors were preserved. The colour scale represents TFCE-corrected p-values, with darker colours indicating smaller p-values. Parcels with p ≥ .05 are transparent. **a.** Unique contribution of semantic root and grammatical function interaction in the Croft interactive model. **b.** Unique contribution of prototypicality in the Croft additive model.

Following the full interaction analysis, we asked a more targeted question: whether neural responses were sensitive to the binary distinction between prototypical and non-prototypical root-function mappings. We encoded a binary predictor of prototypicality as a theoretically motivated summary of root-function mapping structure to replace the root function interaction predictor, and then compared the differences in the explained variances between the true-label model and the permuted model. We found significant contributions of prototypicality as a predictor in subject 1 and subject 3, but it was absent in the subject 2 dataset after multiple-comparisons correction (Figure 3b).

### Prototypical combination coefficients differ across parcels and time

Moving beyond establishing the independent contributions of semantic root and grammatical function to neural prediction, we next tested another important claim in Croft’s framework, that the three prototypical combinations of semantic root and grammatical function, namely, object × reference, action × predication, and property × modification, define prototypical nouns, verbs and adjectives. Rather than testing whether these categories made further overall predictive contributions, we asked how their associated responses differed over the time course of word processing and when these differences emerged. We therefore directly compared their time-resolved coefficients by reconstructing the coefficients for the three prototypes and testing all three pairwise contrasts separately within each participant.

The contrast between object × reference (prototypical noun) and action × predication (prototypical verb) produced the most consistent result (Figure 4). Significant clusters occurred in 13 parcels for subject 1, 11 parcels for subject 2 and 18 parcels for subject 3, most consistently in the left temporal cortex and with some involvement in the right temporal cortex in some of the subjects. 3 parcels were significant to the contrast between the prototypical nouns and verbs across all three datasets, in the left superior temporal cortex. In general, the temporal contrast demonstrated a pattern with early differences around 200ms to 400ms and then later differences around 600ms to 900ms.

**Figure 4.**
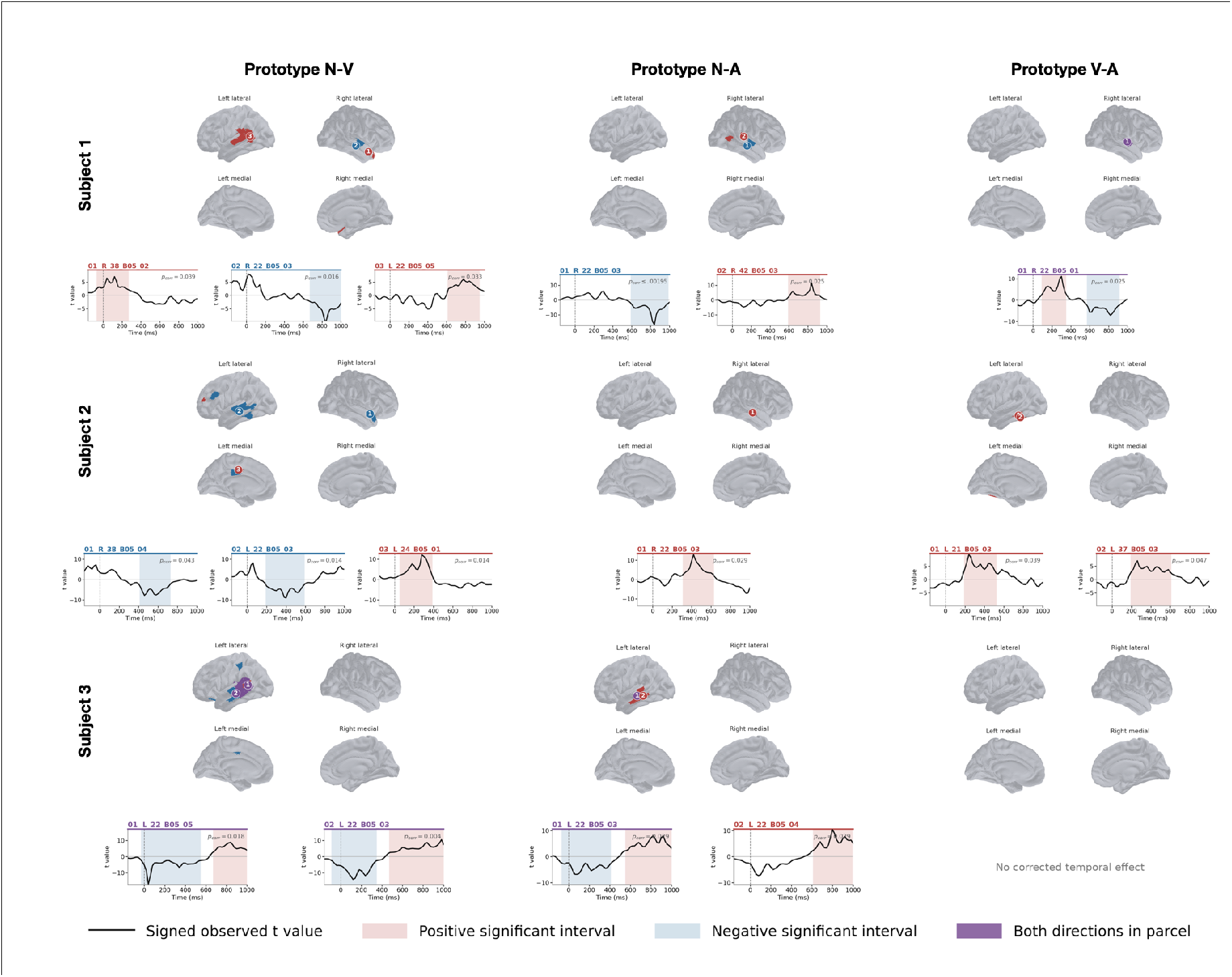
Pairwise comparisons between the coefficients of the three prototypical root-function mappings. Rows correspond to the three participant datasets, and columns show the contrasts object × reference (prototype N) minus action × predication (prototype V), object × reference (prototype N) minus property × modification (prototype A), and action × predication (prototype V) minus property × modification (prototype A). For each contrast, the cortical surface plots show all parcels containing at least one significant temporal for a specific prototype contrast. Red indicates a positive coefficient difference, such that the first prototypical cell had a larger coefficient than the second; blue indicates a negative difference; and purple indicates that significant differences in both directions occurred at different latencies. The time-course panels show the t-values for the significance of the contrasts. Two or three parcels were selected to represent the patterns for each subject and each contrast when more than three parcels are significant for the corresponding contrast.

The other two prototypes’ contrasts were less consistent across datasets. The contrasts between prototypical nouns and prototypical adjectives were significant in 4 parcels for subject 1, 1 parcel for subject 2 and 4 parcels for subject 3, with subject 1 and subject 2 in the right temporal cortex while subject 3 in the left temporal cortex. The significant time windows for the contrast between prototypical nouns and adjectives differ among the three datasets, with a late 600-1000ms time window in subject 1and 3 and an earlier 300-600ms time window in subject 2. The contrasts between prototypical verbs and adjectives were only significant in subject 1 and 2, with 1 parcel in the right superior temporal cortex in subject 1 around 100-350ms, and 2 parcels in the left inferior temporal cortex in subject 2 around 200-450ms.

## Discussion

This study investigated when and where MEG responses during natural language comprehension are sensitive to the linguistic dimensions underlying the major PoS categories of nouns, verbs, and adjectives. Croft’s theory-driven framework (2000, 2001) was operationalized as predictors in time-resolved encoding models. Our results provided neurobiological support for key distinctions in Croft’s framework of content word classes. Croft’s framework distinguishes the semantic root types of words from their grammatical function in the context. For major word classes (nouns, verbs and adjectives), three semantic root types can be distinguished: object, action and property, and three grammatical functions can be defined: reference, predication and modification. And then the combinations between a specific semantic root and a specific grammatical function, though particular pairings can be more typical than others, jointly define word classes. This construction provides a principled explanation for why a word class label can encompass several sources of neural variation. In the present study, the Croft additive model based on the decomposed labels modestly outperformed the model based on conventional core PoS labels in all three subjects. Thus, separating semantic root from grammatical function preserves the word-category information relevant to neural responses while making its internal structure explicit.

### Semantic root and grammatical function as dimensions of word class

The analysis of the unique contribution of each individual predictor provides evidence supporting that semantic root and grammatical function dimensions both matter independently. The conditional tests further demonstrate that grammatical function remained informative in all three participants after controlling for the semantic root types during permutation. Semantic root also retained a conditional contribution in two out of three datasets. The functional contribution therefore cannot be explained entirely by the tendency for particular semantic types to occur in particular grammatical functions. Conversely, the conditional root effects indicate that grammatical function does not exhaust the information captured by semantic type. These results support treating the two dimensions as distinguishable predictors.

Previous studies have extensively investigated the respective contributions of semantic meaning and grammatical class to neural differences between nouns and verbs. Vigliocco et al. (2006) found that neural responses during single-word processing were primarily influenced by sensory and motor meaning, with no significant effect of grammatical class. Similarly, Siri et al. (2008) found no consistent grammatical-class-specific activation when participants named the same events using nominal and verbal forms, with the observed differences in left inferior frontal activity instead reflecting processing demands. On the other hand, Peelen et al. (2012) provided evidence that grammatical class related neural differences cannot always be reduced to semantic distinctions. They found that verb-selective regions in the left lateral temporal cortex responded more strongly to verbs than to semantically matched nouns, with comparable verb selectivity for both event and state meanings. These studies suggest that semantic and grammatical information can make distinguishable contributions to neural responses, although their effects may vary with the linguistic properties and processing contexts being examined. In a systematic review, Vigliocco et al. (2011) discussed different theoretical accounts of grammatical class, including lexicalist, combinatorial, and emergentist views. While lexicalist accounts assume that grammatical class information is specified in lexical representations, combinatorial accounts emphasize grammatical operations during sentence processing. The emergentist view instead proposes that word-class distinctions arise from semantic, pragmatic, and distributional information rather than grammatical class being an independent organizing principle. Our findings are compatible with the emergentist view, showing that semantic root type and grammatical function contribute distinguishable predictive information to neural responses, without requiring word classes to be represented as independent, unitary categories.

Our study extends previous findings in two ways. First, previous research has largely focused on the noun-verb distinction, whereas our study extends the investigation to a broader range of core content words, including adjectives. Second, while previous studies primarily relied on controlled experimental contrasts to separate semantic and grammatical information, our study examined their respective contributions within the same neural encoding model during continuous naturalistic language comprehension. This approach allowed us to assess the contributions of these dimensions as words are processed in an ongoing linguistic context, without requiring explicit tasks. The conditional permutation analyses further allowed us to assess the predictive information carried by each dimension while accounting for its statistical association with the other.

### Cerebral sensitivity to the combinations of semantic root and grammatical function

A purely additive description of semantic root and grammatical function treats each semantic type as contributing in the same way across grammatical functions, and each function as contributing in the same way across semantic types. However, we found that their interaction also provided significant predictive information for neural responses during naturalistic language comprehension. The interaction effects suggest that neural responses are sensitive not only to semantic and grammatical information individually, but also to their specific combinations. A further claim by Croft about the internal organization between semantic roots and grammatical functions is that three mappings, object × reference, action × predication, and property × modification, represent the prototypical definition of nouns, verbs and adjectives. The rest of the combinations often require additional linguistic markings or have more restricted possibilities in natural language occurrence (Croft, 2001, 2022). The importance of the three prototypes for understanding the neural organization of word classes has also been emphasized by Kemmerer (2014), particularly in relation to the prototypical associations between object reference and nouns, and action predication and verbs.

To examine whether this theoretically driven distinction is reflected in neural responses, we included prototypicality (prototypical vs. nonprototypical) as an additional predictor in the Croft additive model. We found that prototypicality contributed significantly to neural prediction in two of the three individual datasets, beyond the individual contributions of semantic root and grammatical function. This finding provides initial neurobiological support for Croft’s proposed prototypical organization of word classes, suggesting that neural responses are sensitive not only to specific root–function combinations, but also to their prototypicality. Interestingly, while the full root × function interaction showed significant contributions across all three participants, prototypicality was significant in only two. This difference raises the possibility that neural sensitivity to root–function combinations involves more detailed distinctions than a single prototypical-nonprototypical contrast can capture.

### Distributed and partially overlapping cortical sensitivity to word-class information

Spatially, our results revealed distributed and partially overlapping cortical patterns for semantic root, grammatical function, and their interaction. Semantic root effects were predominantly left-lateralized, particularly in temporal and frontal regions, consistent with previous findings of left-hemisphere involvement in semantic processing (Vigliocco et al., 2006). Grammatical function showed a broader and more bilateral distribution, whereas interaction effects were concentrated mainly in left temporal and temporoparietal regions. Importantly, seven parcels showed significant contributions from all three predictors across all participants. These parcels were located exclusively in the left temporal cortex, including four in the superior temporal gyrus (BA22), two in the transverse temporal gyrus (BA42), and one in the middle temporal gyrus (BA21). The concentration of these shared effects in temporal regions is consistent with the established role of the superior and middle temporal cortices in speech comprehension and lexical-semantic processing (Hickok & Poeppel, 2007). Previous studies have also shown that superior temporal regions are involved in both semantic and syntactic aspects of sentence processing (Friederici et al., 2003). Our findings extend this perspective by showing that the same cortical parcels carry predictive information about semantic root type, grammatical function, and their specific combinations. These regions may therefore represent important sites of convergence for multiple dimensions of word-class-related information during naturalistic comprehension.

More broadly, this partially shared distribution is consistent with previous evidence against a strict anatomical separation of word classes and their underlying linguistic properties. Crepaldi et al. (2013) reported closely situated noun- and verb-related responses within a distributed fronto-temporo-parietal network, while Bekemeier et al. (2026) showed distributed and context-dependent word-class encoding during language comprehension. Related evidence from Shain et al. (2024) also demonstrated overlapping sensitivity to semantic and syntactic information throughout the language network. These findings support a multidimensional account of word-class processing within a distributed but partly shared cortical network, with the left temporal cortex emerging as a particularly consistent site of sensitivity to semantic, grammatical, and combinational information.

### The spatiotemporal differentiation of prototypical word classes in the brain

To examine whether, when, and where the three prototypical word classes exhibit distinct neural response dynamics, we compared their time-resolved coefficients through pairwise contrasts. These analyses revealed a consistent noun–verb distinction across all three participants, with significant effects concentrated in the left perisylvian cortex (BA22, BA41, and BA42). Although the time windows surviving cluster-mass correction differed across subjects, with a late effect in subject 1, an early effect in subject 2, and both early and late effects in subject 3, the unthresholded statistics in this region consistently followed an early-negative, late-positive pattern. The variation in corrected intervals may therefore reflect differences in the statistical threshold rather than qualitatively different responses. In contrast, noun–adjective differences were weaker and inconsistent in direction and latency, while verb–adjective differences were absent in subject 3.

The reproducibility of the noun–verb contrast is consistent with evidence that the clearest neural distinction between word classes arises for their semantic and functional prototypes, namely object-referring nouns and action-predicating verbs (Vigliocco et al., 2011). The left perisylvian distribution of this effect is consistent with previous evidence that nouns and verbs are differentiated within a shared language network rather than processed in separate category-specific regions (Crepaldi et al., 2013). The weaker and less consistent adjective-related contrasts may reflect the intermediate position of adjectives in Croft’s conceptual space, in which the adjective prototype combines the intermediate semantic class of properties with the intermediate function of modification, while the noun and verb prototypes occupy the contrasting object–reference and action–predication endpoints (Croft, 1991, 2001). Adjectives may therefore share representational properties with both endpoint prototypes rather than forming an equally opposed category, producing less stable pairwise differentiation across participants and time.

### Limitations and future directions

The intensive sampling provided substantial data for within-participant analyses, but the study included only three participants. The findings should therefore be interpreted as evidence from individual datasets rather than population-level estimates, and their generalizability requires testing in a larger sample. The naturalistic design also limits the separation of linguistic dimensions: semantic root types, grammatical functions, and their combinations occur with unequal frequencies. The naturalistic approach indeed reflects how language is ordinarily used, however, unequal numbers of observations across prototypical cells may affect the precision of coefficient estimates and the sensitivity of pairwise comparisons. The present findings should therefore be interpreted alongside evidence from well-controlled experiments that systematically manipulate linguistic properties.

Croft’s framework is motivated by the hunt for cross-linguistic universality, which offers a direction for future research. Extending this decompositional approach to other typologically different languages would allow researchers to examine whether the neural contributions of semantic root, grammatical function, and their combinations generalize across languages, and how these contributions vary with language-specific patterns of grammatical encoding.

### Conclusions

By approaching parts of speech from a dimensional rather than a unitary categorical perspective under Croft’s framework, this study shows that semantic root and grammatical function make independent contributions to MEG responses during narrative comprehension, while their combinations provide additional predictive information in specific cortical regions. Semantic root type and grammatical function each contributed unique predictive information, while their interaction contributed beyond their additive effects, showing that PoS-related sensitivity cannot be attributed solely to semantic meaning or grammatical role. The interaction was reliable across all three datasets, whereas binary prototypicality contributed in two, suggesting sensitivity to detailed root-function mappings rather than only a prototypical–nonprototypical distinction. The reproducible temporal differentiation between prototypical nouns and verbs reveals how they emerge from particular root-function mappings and unfold during word processing; prototype adjective-related differences were less stable, potentially reflecting the intermediate position of the property–modification mapping in Croft’s conceptual space between the object–reference and action–predication prototypes. These findings support a decompositional account in which PoS-related effects arise from distributed, partially overlapping sensitivity to semantic root type, grammatical function, and their combinations.

## Methods

### Participants

The dataset comprises three right-handed native speakers of English, who spoke UK, US, or South African English. The three participants, one female and 2 male, aged 35,30, and 28, all had normal or corrected-to-normal vision and reported no history of neurological, developmental, or language disorders. Participants provided written informed consent, including consent for anonymized data sharing. The original study was approved by the local Committee on Research Involving Human Subjects in the Arnhem-Nijmegen region and was conducted in accordance with the Declaration of Helsinki.

### Procedure

Participants listened to continuous narrative speech of ten stories from The Adventures of Sherlock Holmes by Arthur Conan Doyle, narrated by David Clarke. Each participant completed ten one-hour MEG sessions on separate days, with one story presented per session. Each story was divided at meaningful narrative boundaries into six to eight runs of approximately 10 min. The order of stories and runs was held constant across participants. Participants were instructed to listen attentively for comprehension and to minimize movement. Participants wore individualized MEG-compatible headcasts to reduce head movement. After each run, they answered comprehension questions and rated the informativeness and absorption of the preceding passage on seven-point scales. Short breaks were permitted between runs. After all MEG sessions, structural MRI was acquired with a 3T Siemens MAGNETOM Skyra scanner and was used for participant-specific anatomy and source reconstruction.

### Stimulus annotation

Word and phoneme onset times were extracted for each word and phoneme by performing the forced alignment between the audiobook recordings and the corresponding story transcripts using the Penn Forced Aligner (P2FA). The resulting word and phoneme intervals were manually checked and corrected when an annotated pronunciation did not match the pronunciation dictionary used by the phoneme model.

Acoustic-level control variables include broadband-envelope variance, phoneme surprisal, uniqueness point, pitch and spectral variation. Phoneme identity was taken from the ARPABET transcription produced by forced alignment. Phoneme surprisal was calculated as the negative log probability of the observed phoneme. Broadband-envelope variance was computed over the duration of each phoneme to capture rapid changes in envelope amplitude. A uniqueness point marked the phoneme at which a word became uniquely identifiable within the pronunciation hierarchy. Mean pitch was extracted for each phoneme with Praat. Pitch was set to zero for voiceless phonemes or intervals in which no pitch could be identified. Spectral variation was represented by a log-mel spectrogram with eight bands, summarized as the mean amplitude within each band over the duration of each phoneme.

Word-level control variables were defined at word or content word onsets. At each word onset, lexical surprisal and lexical frequency were encoded. Lexical surprisal was the negative log conditional probability of the observed word under GPT-2 XL (Radford et al., 2019). Lexical frequency was presented as an inverse-frequency measure, computed as the negative log probability of the word based on its corpus frequency in SUBTLEX. At each content word onset, semantic mismatch quantified the cosine distance between the embedding of the presented word and a predicted semantic vector obtained by probability-weighting the GloVe embeddings (Pennington et al., 2014) of candidate words in the GPT-2 distribution. Contextual dissimilarity controlled for bottom-up semantic association or integration difficulty, computed as the cosine distance between its semantic embedding and the average embedding of the preceding context words.

More importantly, each content word was assigned one semantic root category (object, action, or property) and one grammatical function category (reference, predication, or modification). Three linguists manually annotated each content word for semantic root and grammatical function, following Croft’s framework (2001, 2022). The annotations were subsequently double-checked. Within the 3 × 3 interaction produced by the combination of semantic categories and grammatical categories, object × reference, action × predication, and property × modification were marked as prototypical, while the six off-diagonal combinations were defined as non-prototypical. Root and function categories were each represented by two contrast-coded predictors, action and object, and predication and reference, with property and modification serving as their respective reference categories. Then the four pairwise products captured their interactions.

### Preprocessing

MEG data were preprocessed in MATLAB using FieldTrip (Oostenveld et al., 2011), following the pipeline released with the dataset. Continuous data were demeaned, high-pass filtered at 0.5 Hz, and notch filtered at 49 Hz -51 Hz, 99 Hz -101 Hz, and 149 Hz -151 Hz. Bad channels and segments containing muscle artifacts or SQUID jumps provided by the public dataset were excluded, and ocular and cardiac artifacts were removed using ICA. Data were subsequently low-pass filtered at 8 Hz. We focused on such slow-evoked responses following the common practice in existing literature using regression ERP or temporal response function (TRF) analysis, especially when the regressors are sparse impulses (e.g., Broderick et al., 2018; Ding et al., 2012; Heilbron et al., 2022). Then the data were downsampled from 1200Hz to 150 Hz. Artifact-contaminated samples were represented as missing values and excluded from further analysis.

### Source reconstruction

Participant-specific MRI-MEG co-registrations, cortical surface reconstructions, volume-conduction models, and leadfields were provided with the dataset. Cortical surfaces were reconstructed with FreeSurfer and downsampled to 7,842 source locations per hemisphere; leadfields were computed using a single-shell volume-conduction model. Source activity was estimated separately for each participant and session using a linearly constrained minimum variance (LCMV) beamformer (Van Veen et al. 1997) implemented in FieldTrip. Source estimates were subsequently parcellated using the 374-parcel refinement of the Conte69/Brodmann atlas, excluding unknown and medial-wall parcels. For each parcel, the first principal component of the constituent source time series was retained as the representative time series per parcel.

### Model construction and comparisons

#### Encoding model estimation

Time-resolved regression models were fitted separately for each participant within a regression ERP framework, following Heilbron et al. (2022). The regression ERP represents predictor values as event-based sparse impulses at word or phoneme onsets and temporally expands those event-based regressors, making each predictor column a series of columns over a range of temporal lags. The models were estimated using ridge regression (regularization parameter λ = 10,000). Each predictor was time-expanded from 150 ms before to 1,000 ms after its event onset. Continuous covariates were z-scored within each run, and categorical predictors retained their specified contrast codes. Model performance was estimated by leave-one-run-out cross-validation: each fold held out one run and trained the model on all remaining runs. For each held-out run and cortical parcel, model performance was quantified by the Pearson correlation (r) and coefficient of determination (R²) between the predicted and observed source time series.

#### Overall performance model comparison

Three models that differed only in how content word category information were constructed and compared. They shared the same base model, including word onsets, lexical surprisal and word frequency at word level, semantic mismatch and contextual dissimilarity at content word level, and broadband envelope variance, pitch and spectrogram amplitudes at phoneme level. The conventional spaCy PoS model additionally included the content word PoS labels generated by spaCy at the content word level. The Croft additive model contained semantic root and grammatical function labels at the content word level. The Croft interaction model contained semantic root and grammatical function labels and their interaction labels at the content word level. We made three prediction performance comparisons: Croft additive versus spaCy PoS, Croft interaction versus spaCy PoS, and Croft interaction versus Croft additive.

#### Models for predictor contribution analyses

We identify the unique contribution of each predictor as the differences in the explained variance across cortical parcels between a true model and a permuted model. The permutation for each categorical control model used one reproducible, prespecified randomization. A true model and a permuted model had the same regressor structure and the same time-lagged design structure for matched model complexity control. In the permuted model, only the values of the predictor being tested were randomized. Three reference model families were fitted. The additive model (MA) contained the predictors in the base model, together with the semantic root predictor and grammatical function predictor at the content word level. The interaction model (MI) contained the predictors in the base model, together with the semantic root predictor, grammatical function predictor, and their interaction. The prototypicality model (MC) contained the common controls, both main semantic root and grammatical function effects, and a single binary prototypicality predictor. Contributions were always evaluated within the same model family.

#### Root and function contributions

Root and function were tested in the MA model family. To estimate the general contribution of root, root labels were shuffled across content word within each run and the two root contrast columns were recomputed. Function was tested analogously by shuffling function labels within each run while retaining the true root columns. We also performed conditional tests in which we preserved the observed level of the other factor: root labels were shuffled separately within each function level and run, and function labels were shuffled separately within each root level and run. These comparisons tested whether neural responses are still sensitive to each dimension after controlling the other dimension.

#### Interaction and control-predictor contributions

The root*function interaction was tested in the MI model family. In the interaction-permuted model, the interaction columns were recomputed from independently shuffled root and function labels within each run. This preserved the within-run marginal frequencies of both predictor labels while disrupting their observed pairings. The same interaction reference model was also used to test the unique contribution of control predictors, including lexical surprisal, unigram surprisal, and an acoustic block comprising broadband-envelope variance, pitch, and the eight spectrographic-band amplitudes. For these controls, the selected predictor or predictor block was also permuted within each run.

#### Prototypicality contribution

Prototypicality was coded as +0.5 for the three prototypical root–function mappings and −0.5 for the six non-prototypical mappings. It was tested in the MC model family by comparing the true model with the permuted model in which function assignments were shuffled within each root level and each run, and the prototypicality value was then recomputed.

#### Prototypical-cell coefficient estimation and pairwise contrasts

To characterize how the fitted coefficients differentiated the three prototypical root-function mappings, we used each session as an individual observation to conduct pairwise comparison of the coefficients. For each participant, we independently re-estimated the true interaction model (MI) within each session, pooling all runs within a session. These session-wise fits used the same predictor set, sum coding, fixed ridge penalty, source-space data, 0.5–8 Hz frequency range, and time lag window as the corresponding all-session MI model, fitting ten non-overlapping session-level coefficient estimates for each participant. From each fit, we reconstructed the coefficient time course of the three prototypical cells from the explicit sum-coded coefficients for semantic root, grammatical function and their interaction. At every parcel and lag, we then formed the three pairwise contrasts: Object × Reference – Action × Predication, Object × Reference − Property × Modification, and Action × Predication − Property × Modification.

### Statistical testing

#### Analysis level

All statistical analyses were conducted separately for each participant, treating the three participants as three individual long-duration datasets. The held-out runs provided the observations for within-dataset inference. Data were not pooled across participants.

#### Overall model performance

For every cross-validation fold, parcel-wise prediction correlations were averaged across the 100 predefined language network parcels (see in Supplementary material). For each of the three planned comparisons, the paired fold-wise difference in mean correlation was tested against zero. A two-sided one-sample t-test was used when the distribution of differences passed the normality and outlier checks; otherwise, a two-sided Wilcoxon signed-rank test was used. This procedure selected a t-test for participant 1 and Wilcoxon tests for participants 2 and 3. The three p-values were Holm-corrected within each participant.

#### Unique predictor contributions

For each predictor contribution comparison, unique predictive contribution was quantified at every parcel and held-out fold as ΔR² = R²_true_ − R²_permuted_, where the true and permuted scores came from matched models within the same family. Positive values therefore indicate higher held-out explained variance for the true labels than for their matched randomized control.

#### Spatial inference

For each participant and model comparison, fold-wise ΔR² values were tested against zero at each parcel with a one-sample t statistic. Multiple comparisons across the 370 cortical parcels were controlled using a two-sided sign-flip permutation test with threshold-free cluster enhancement (TFCE) (Smith and Nichols, 2009), incorporating the atlas-based adjacency between parcels. We used 9,999 inferential permutations and a TFCE-corrected significance threshold of p < .05.

#### Prototypical-cell coefficient contrasts

Within each participant, at every parcel and time point from -150 to 1000 ms, each of the three prototypical-cell coefficient differences was tested against zero using a one-sample t-test across the ten session estimates. The null distribution was obtained by exact two-sided sign flipping^1^. The coefficient inference used a temporal cluster-mass test (Maris & Oostenveld, 2007). Temporal clusters were formed separately within each contrast and parcel from consecutive samples that exceeded the two-sided p < .05 cluster-forming threshold with 9 degrees of freedom. Cluster mass was the sum of the absolute t values in the cluster. For each sign pattern, the largest cluster mass across all three contrasts and all parcels was entered into a joint maximum null distribution. Corrected cluster p-values were obtained by comparing each observed mass with this distribution, controlling the family-wise error rate across contrasts, parcels, and the tested temporal window at p < .05.

#### Data availability

The 10-hour within-participant MEG dataset was published by Armeni et al. (2022), available at https://data.ru.nl/collections/di/dccn/DSC_3011085.05_995. The language materials, annotations and code used to produce the results in this manuscript are available at https://github.com/proposaluzh/sherlock_trf.

## Acknowledgements

This work is supported by the Swiss National Science Foundation (Sinergia grant No. CRSII5_209413). We acknowledge the authors, Armeni et al. (2022), for the publicly available dataset used in this study and Heilbron et al. (2022) for making their analysis pipeline publicly available. The authors are grateful to Elizabeth Clara Wagner for assistance with the linguistic annotation.

## Footnotes

1 Since there are 10 sessions as observations, all 2⁹ = 512 unique sign patterns were enumerated.

## References

Armeni, K., Güçlü, U., van Gerven, M., & Schoffelen, J.-M. (2022). A 10-hour within-participant magnetoencephalography narrative dataset to test models of language comprehension. Scientific Data, 9, 278. 10.1038/s41597-022-01382-7

Bekemeier, N., Hundt, M., Huang, Z., Djordjijevic, M., & Hervais-Adelman, A. (2026). Cerebral encoding of word classes is distributed and context-dependent [Preprint]. bioRxiv. 10.64898/2026.07.22.739973

Broderick, M. P., Anderson, A. J., Di Liberto, G. M., Crosse, M. J., & Lalor, E. C. (2018). Electrophysiological correlates of semantic dissimilarity reflect the comprehension of natural, narrative speech. Current Biology, 28(5), 803–809.e3. 10.1016/j.cub.2018.01.080

Crepaldi, D., Berlingeri, M., Cattinelli, I., Borghese, N. A., Luzzatti, C., & Paulesu, E. (2013). Clustering the lexicon in the brain: A meta-analysis of the neurofunctional evidence on noun and verb processing. Frontiers in Human Neuroscience, 7, 303. 10.3389/fnhum.2013.00303

Croft, W. (1991). Syntactic categories and grammatical relations: The cognitive organization of information. University of Chicago Press.

Croft, W. (2000). Parts of speech as language universals and as language-particular categories. In P. M. Vogel & B. Comrie (Eds.), Approaches to the typology of word classes (pp. 65–102). Mouton de Gruyter. 10.1515/9783110806120.65

Croft, W. (2001). Radical construction grammar: Syntactic theory in typological perspective. OUP Oxford.

Croft, W. (2002). Typology and Universals (2 ed.). Cambridge University Press. DOI: 10.1017/CBO9780511840579

Croft, W. (2022). Word classes in radical construction grammar. In: Oxford handbook of word classes. Oxford University Press.

Crosse, M. J., Di Liberto, G. M., Bednar, A., & Lalor, E. C. (2016). The multivariate temporal response function (mTRF) toolbox: A MATLAB toolbox for relating neural signals to continuous stimuli. Frontiers in Human Neuroscience, 10, 604. 10.3389/fnhum.2016.00604

Damasio, A. R., & Tranel, D. (1993). Nouns and verbs are retrieved with differently distributed neural systems. Proceedings of the National Academy of Sciences, 90(11), 4957–4960. 10.1073/pnas.90.11.4957

Ding, N., & Simon, J. Z. (2012). Neural coding of continuous speech in auditory cortex during monaural and dichotic listening. Journal of Neurophysiology, 107(1), 78–89. 10.1152/jn.00297.2011

Federmeier, K. D., Segal, J. B., Lombrozo, T., & Kutas, M. (2000). Brain responses to nouns, verbs and class-ambiguous words in context. Brain, 123(12), 2552–2566. 10.1093/brain/123.12.2552

Friederici, A. D., Rüschemeyer, S.-A., Hahne, A., & Fiebach, C. J. (2003). The role of left inferior frontal and superior temporal cortex in sentence comprehension: Localizing syntactic and semantic processes. Cerebral Cortex, 13(2), 170–177. 10.1093/cercor/13.2.170

Heilbron, M., Armeni, K., Schoffelen, J.-M., Hagoort, P., & de Lange, F. P. (2022). A hierarchy of linguistic predictions during natural language comprehension. Proceedings of the National Academy of Sciences, 119(32), e2201968119. 10.1073/pnas.2201968119

Hickok, G., & Poeppel, D. (2007). The cortical organization of speech processing. Nature Reviews Neuroscience, 8(5), 393–402. 10.1038/nrn2113

Honnibal, M., Montani, I., Van Landeghem, S., & Boyd, A. (2020). spaCy: Industrial-strength natural language processing in Python [Computer software]. Zenodo. 10.5281/zenodo.1212303

Kemmerer, D. (2014). Word classes in the brain: Implications of linguistic typology for cognitive neuroscience. Cortex, 58, 27–51. 10.1016/j.cortex.2014.05.004

Maris, E., & Oostenveld, R. (2007). Nonparametric statistical testing of EEG- and MEG-data. Journal of Neuroscience Methods, 164(1), 177–190. 10.1016/j.jneumeth.2007.03.024

Oostenveld, R., Fries, P., Maris, E., & Schoffelen, J.-M. (2011). FieldTrip: Open source software for advanced analysis of MEG, EEG, and invasive electrophysiological data. Computational Intelligence and Neuroscience, 2011, Article 156869. 10.1155/2011/156869

Peelen, M. V., Romagno, D., & Caramazza, A. (2012). Independent representations of verbs and actions in left lateral temporal cortex. Journal of Cognitive Neuroscience, 24(10), 2096–2107. 10.1162/jocn_a_00257

Pennington, J., Socher, R., & Manning, C. D. (2014). GloVe: Global vectors for word representation. In Proceedings of the 2014 Conference on Empirical Methods in Natural Language Processing (EMNLP) (pp. 1532–1543). Association for Computational Linguistics. 10.3115/v1/D14-1162

Pulvermüller, F., Lutzenberger, W., & Preissl, H. (1999). Nouns and verbs in the intact brain: Evidence from event-related potentials and high-frequency cortical responses. Cerebral Cortex, 9(5), 497–506. 10.1093/cercor/9.5.497

Radford, A., Wu, J., Child, R., Luan, D., Amodei, D., & Sutskever, I. (2019). Language models are unsupervised multitask learners. OpenAI. https://cdn.openai.com/better-language-models/language_models_are_unsupervised_multitask_learners.pdf

Shain, C., Kean, H., Casto, C., Lipkin, B., Affourtit, J., Siegelman, M., Mollica, F., & Fedorenko, E. (2024). Distributed sensitivity to syntax and semantics throughout the language network. Journal of Cognitive Neuroscience, 36(7), 1427–1471. 10.1162/jocn_a_02164

Shapiro, K. A., Moo, L. R., & Caramazza, A. (2006). Cortical signatures of noun and verb production. Proceedings of the National Academy of Sciences, 103(5), 1644–1649. 10.1073/pnas.0504142103

Siri, S., Tettamanti, M., Cappa, S. F., Della Rosa, P., Saccuman, C., Scifo, P., & Vigliocco, G. (2008). The neural substrate of naming events: Effects of processing demands but not of grammatical class. Cerebral Cortex, 18(1), 171–177. 10.1093/cercor/bhm043

Smith, N. J., & Kutas, M. (2015a). Regression-based estimation of ERP waveforms: I. The rERP framework. Psychophysiology, 52(2), 157–168. 10.1111/psyp.12317

Smith, N. J., & Kutas, M. (2015b). Regression-based estimation of ERP waveforms: II. Nonlinear effects, overlap correction, and practical considerations. Psychophysiology, 52(2), 169–181. 10.1111/psyp.12320

Smith, S. M., & Nichols, T. E. (2009). Threshold-free cluster enhancement: Addressing problems of smoothing, threshold dependence and localisation in cluster inference. NeuroImage, 44(1), 83–98. 10.1016/j.neuroimage.2008.03.061

Van Veen, B. D., van Drongelen, W., Yuchtman, M., & Suzuki, A. (1997). Localization of brain electrical activity via linearly constrained minimum variance spatial filtering. IEEE Transactions on Biomedical Engineering, 44(9), 867–880. 10.1109/10.623056

Vigliocco, G., Warren, J., Siri, S., Arciuli, J., Scott, S., & Wise, R. (2006). The role of semantics and grammatical class in the neural representation of words. Cerebral Cortex, 16(12), 1790–1796. 10.1093/cercor/bhj115

Vigliocco, G., Vinson, D. P., Druks, J., Barber, H., & Cappa, S. F. (2011). Nouns and verbs in the brain: A review of behavioural, electrophysiological, neuropsychological and imaging studies. Neuroscience & Biobehavioral Reviews, 35(3), 407–426. 10.1016/j.neubiorev.2010.04.007

