## Supplemental information for "Decomposing Parts of Speech: Cerebral Encoding of Semantic and Grammatical Dimensions and Their Combinational Structure"

Zirui Huang[1]*, Natalia Bekemeier[1], Alexis Hervais-Adelman [1,2]
[1] Zurich Linguistics Centre, University of Zurich, Zurich, Switzerland
[2] Department of Basic Neurosciences, University of Geneva, Geneva, Switzerland

| | | Grammatical function | | | |
|---|---|---|---|---|---|
| | | Reference | Modification | Predication | Total |
| Semantic Root | Object | 12911 | 884 | 173 | 13968 |
| | Property | 507 | 3847 | 1076 | 5430 |
| | Action | 2433 | 630 | 9792 | 12855 |
| | Total | 15851 | 5361 | 11041 | 32253 |

**Table S1. Distribution of semantic-root and grammatical-function annotations.**

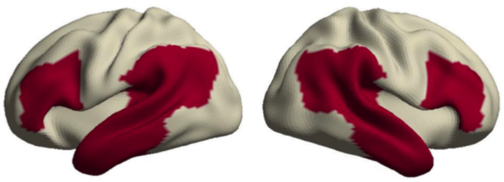

**Figure S1. Language network definition.** Following Heilbron et al. (2022), the language network was broadly defined as the bilateral temporal cortex, temporoparietal junction, inferior frontal gyrus, and dorsolateral prefrontal cortex, comprising 100 of the 370 cortical parcels. This mask was used to summarize overall predictive performance for the model comparisons presented in Figure 1.

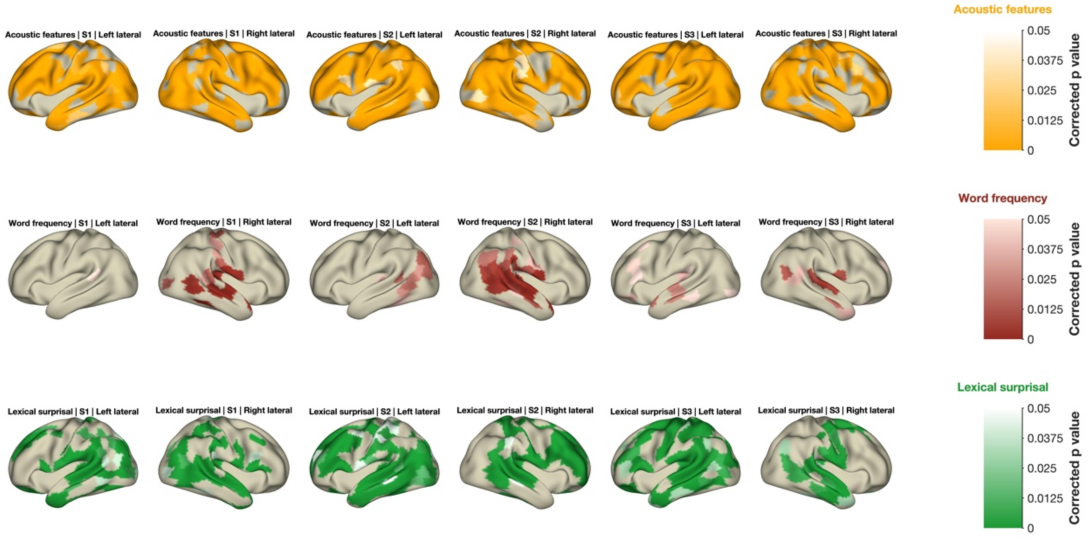


**Figure S2. Unique predictive contributions of the control predictors.** Spatially resolved model comparisons show the unique contributions of acoustic features (top), word frequency (middle), and lexical surprisal (bottom) to predicting MEG responses in each participant. For each predictor or predictor block, the full model was compared with a matched model in which only the corresponding regressors were permuted. Left and right lateral views are shown for participants 1–3. Colours represent TFCE-corrected p-values, with darker colours indicating smaller p-values. Parcels with corrected p ⩾ .05 are transparent.

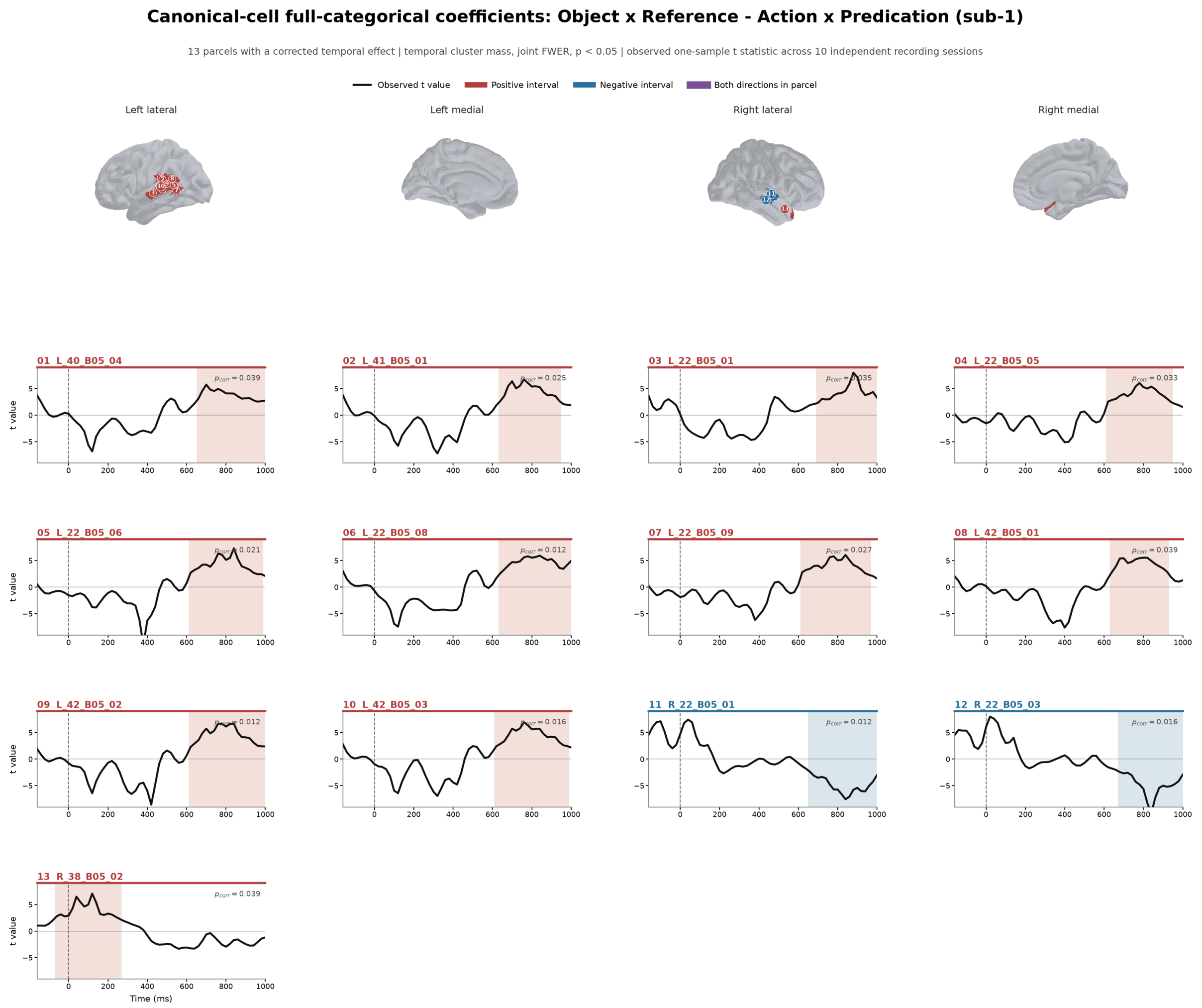


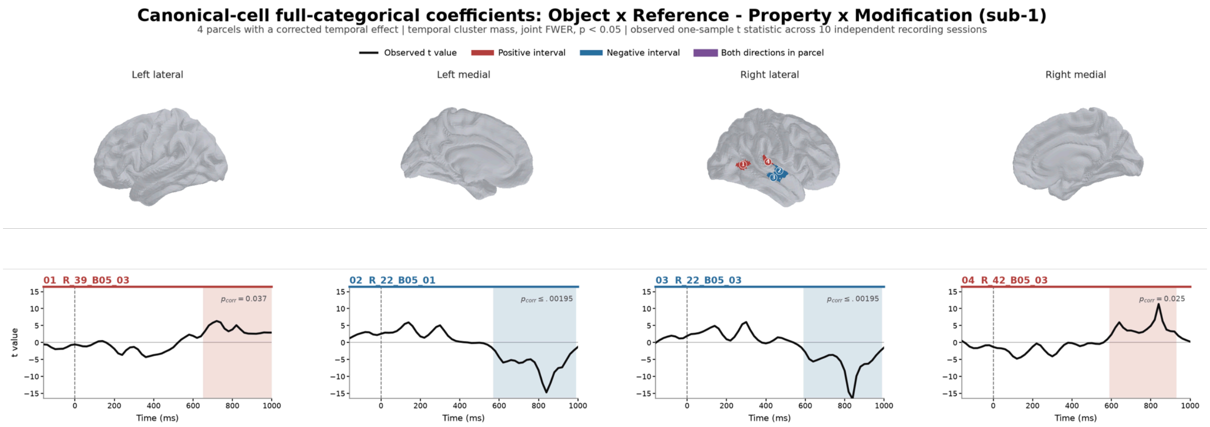

### Canonical-cell full-categorical coefficients: Action x Predication - Property x Modification (sub-1)

1 parcels with a corrected temporal effect | temporal cluster mass, joint FWER, $p < 0.05$ | observed one-sample t statistic across 10 independent recording sessions

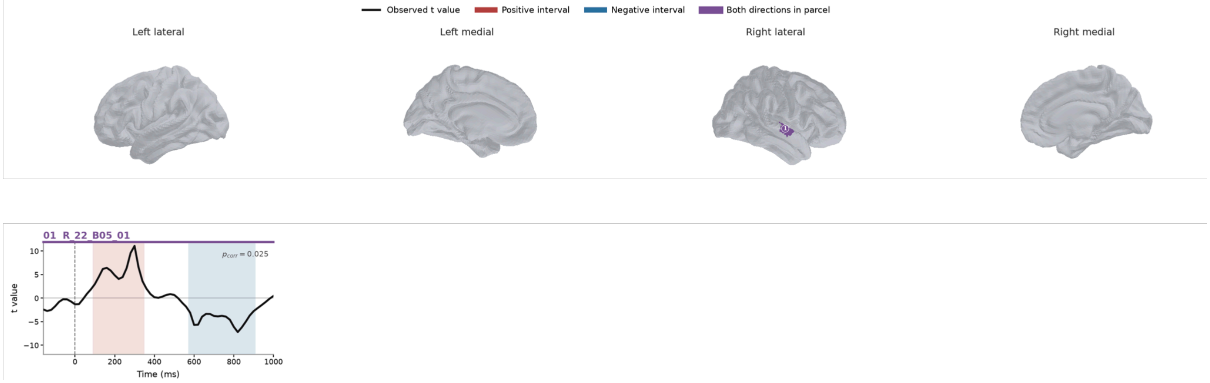


### Canonical-cell full-categorical coefficients: Object x Reference - Action x Predication (sub-2)

11 parcels with a corrected temporal effect | temporal cluster mass, joint FWER, $p < 0.05$ | observed one-sample t statistic across 10 independent recording sessions

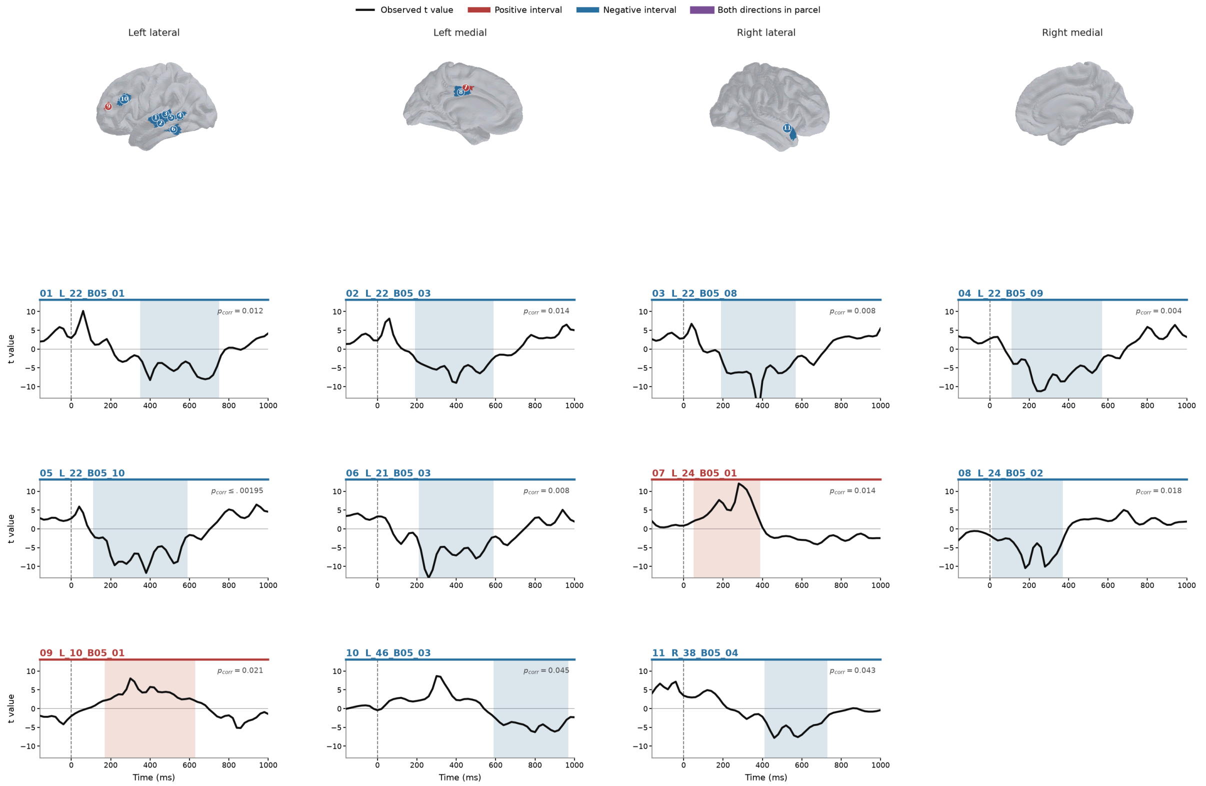

### Canonical-cell full-categorical coefficients: Object x Reference - Property x Modification (sub-2)

1 parcels with a corrected temporal effect | temporal cluster mass, joint FWER, p < 0.05 | observed one-sample t statistic across 10 independent recording sessions

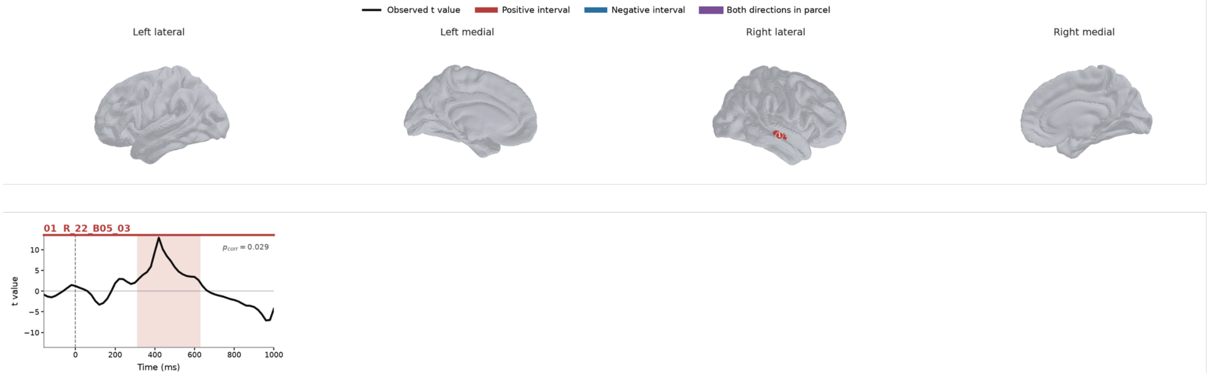


### Canonical-cell full-categorical coefficients: Action x Predication - Property x Modification (sub-2)

2 parcels with a corrected temporal effect | temporal cluster mass, joint FWER, p < 0.05 | observed one-sample t statistic across 10 independent recording sessions

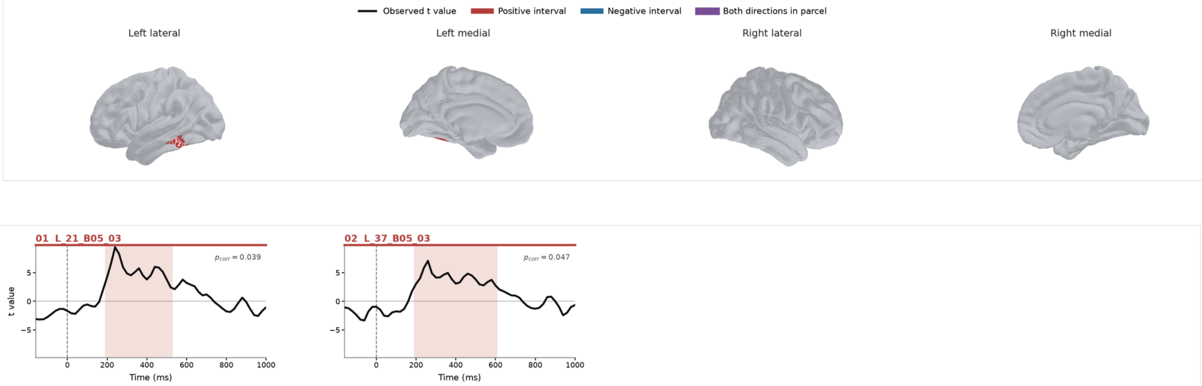

### Canonical-cell full-categorical coefficients: Object x Reference - Action x Predication (sub-3)

18 parcels with a corrected temporal effect | temporal cluster mass, joint FWER, p < 0.05 | observed one-sample t statistic across 10 independent recording sessions

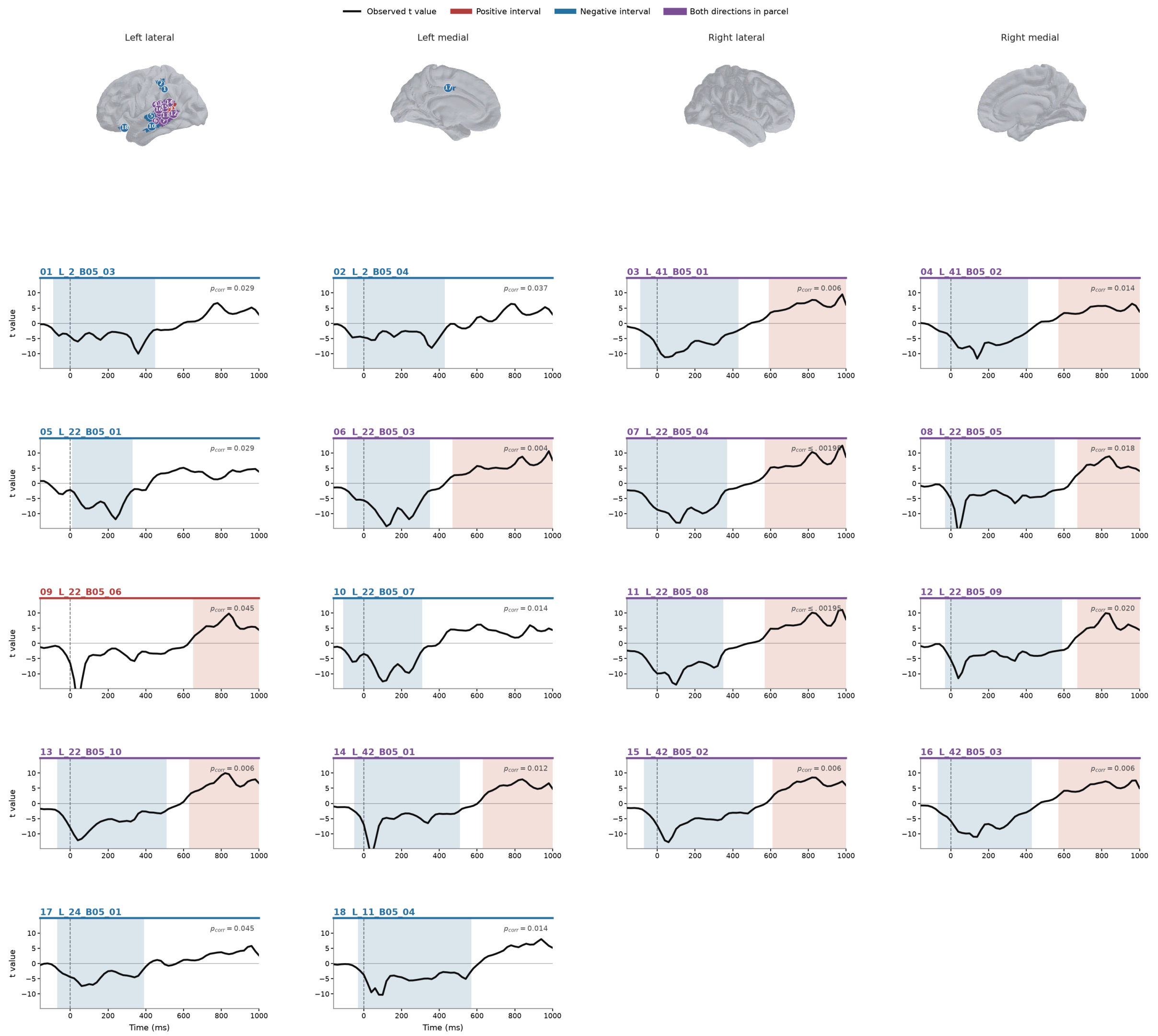


### Canonical-cell full-categorical coefficients: Object x Reference - Property x Modification (sub-3)

4 parcels with a corrected temporal effect | temporal cluster mass, joint FWER, p < 0.05 | observed one-sample t statistic across 10 independent recording sessions

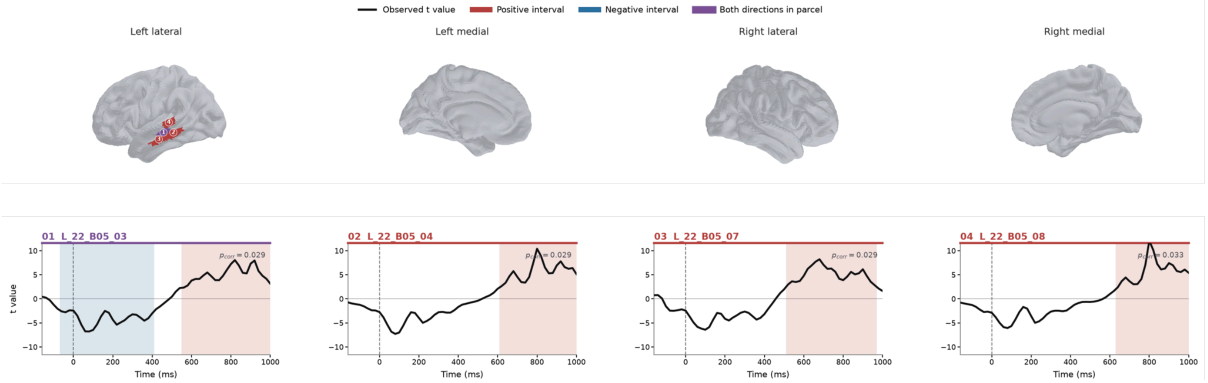

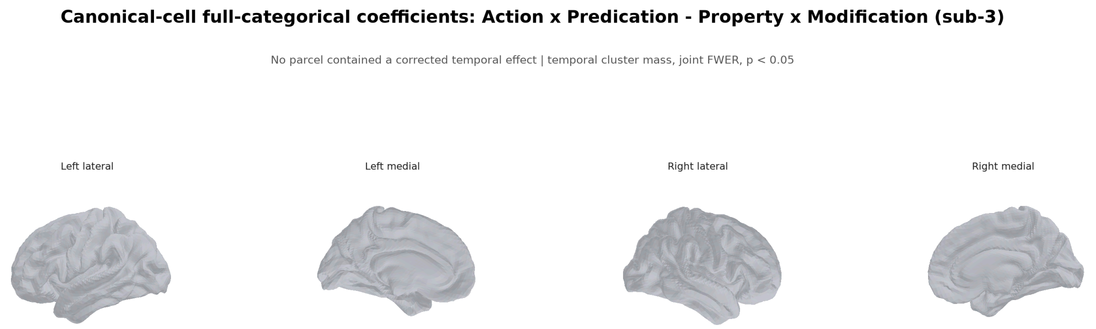


**Figure S3. Complete parcel-wise time courses for the pairwise contrasts between prototypical word classes.** The figure presents all parcels showing a significant difference between the reconstructed coefficients of prototypical nouns (Object × Reference), verbs (Action × Predication), and adjectives (Property × Modification) in each participant. Numbered labels on the lateral and medial cortical maps correspond to the time-course panels. Black lines show the signed observed t-values across the ten session-wise coefficient estimates. Shaded regions indicate significant temporal clusters, with red denoting a positive contrast and blue a negative contrast. Cortical parcels are coloured according to the direction of their significant effects; purple indicates parcels containing significant intervals in both directions. Family-wise error was controlled jointly across the three contrasts, all cortical parcels, and tested time points within each participant (p < .05).
